# Thermoneutral housing does not accelerate metabolic dysfunction-associated fatty liver disease in male or female mice fed a Western diet

**DOI:** 10.1101/2023.01.24.524609

**Authors:** Julia R.C. Nunes, Tyler K.T. Smith, Peyman Ghorbani, Conor O’Dwyer, Natasha A. Trzaskalski, Habiba Dergham, Ciara Pember, Marisa K. Kilgour, Erin E. Mulvihill, Morgan D. Fullerton

## Abstract

**Objective:** Metabolic dysfunction-associated fatty liver disease (MAFLD) represents a growing cause of mortality and morbidity and encompasses a spectrum of liver pathologies. Potential therapeutic targets have been identified and are currently being pre-clinically and clinically tested. However, while dozens of preclinical models have been developed to recapitulate various stages of MAFLD, few achieve fibrosis using an experimental design that mimics human pathogenesis. We sought to clarify whether the combination of thermoneutral (T_N_) housing and consumption of a classical Western diet (WD) would accelerate the onset of MAFLD and progression in male and female mice.

**Methods:** Male and female C57Bl/6J mice were fed a nutrient-matched low-fat control or Western diet (41% Kcal from fat, 43% carbohydrate and 0.2% cholesterol; WD) starting at ∼12 wk of age for a further 16 wk. Mice were divided and housed with littermates at either standard temperature (T_S_; 22°C) or thermoneutral conditions (T_N_; ∼29°C). Mice underwent tests for glucose tolerance, insulin sensitivity and body composition, as well as intestinal permeability. Following tissue harvest, circulating and liver markers of hepatic disease progression toward steatosis and fibrosis were determined.

**Results:** While male mice housed at T_N_ and fed a WD were significantly heavier than T_S_ -housed control animals, no other differences in body weight or composition were observed. WD-fed females housed under T_N_ conditions had higher circulating LDL-cholesterol; however, there were no other significant differences between T_N_ and T_S_ -housing in circulating or hepatic lipid levels. While WD-fed T_N_ males had higher ALT levels, no other differences in markers of liver injury or disease progression were observed. Moreover, females housed at T_N_ conditions and fed a WD remained significantly protected against the induction of fibrosis compared to male counterparts. Interestingly, sex-specific differences were observed in markers of glucose and insulin tolerance, where T_N_ housing and WD feeding resulted in hyperglycemia and impaired insulin responsiveness in both sexes, but glucose intolerance only in male mice.

**Conclusions:** While T_N_ housing has been demonstrated to exacerbate high fat-induced hepatic steatosis and inflammation in male and female mice, coupling T_N_ housing with a WD for 16 wk was not sufficient to augment fatty liver progression in male or female mice.

**Highlights:**

1. Thermoneutral housing and Western diet feeding does not progress to NASH
2. Female mice are not more susceptible to obesity induced fatty liver under these conditions
3. Temperature and diet had sex-specific effects on glucose tolerance and insulin sensitivity

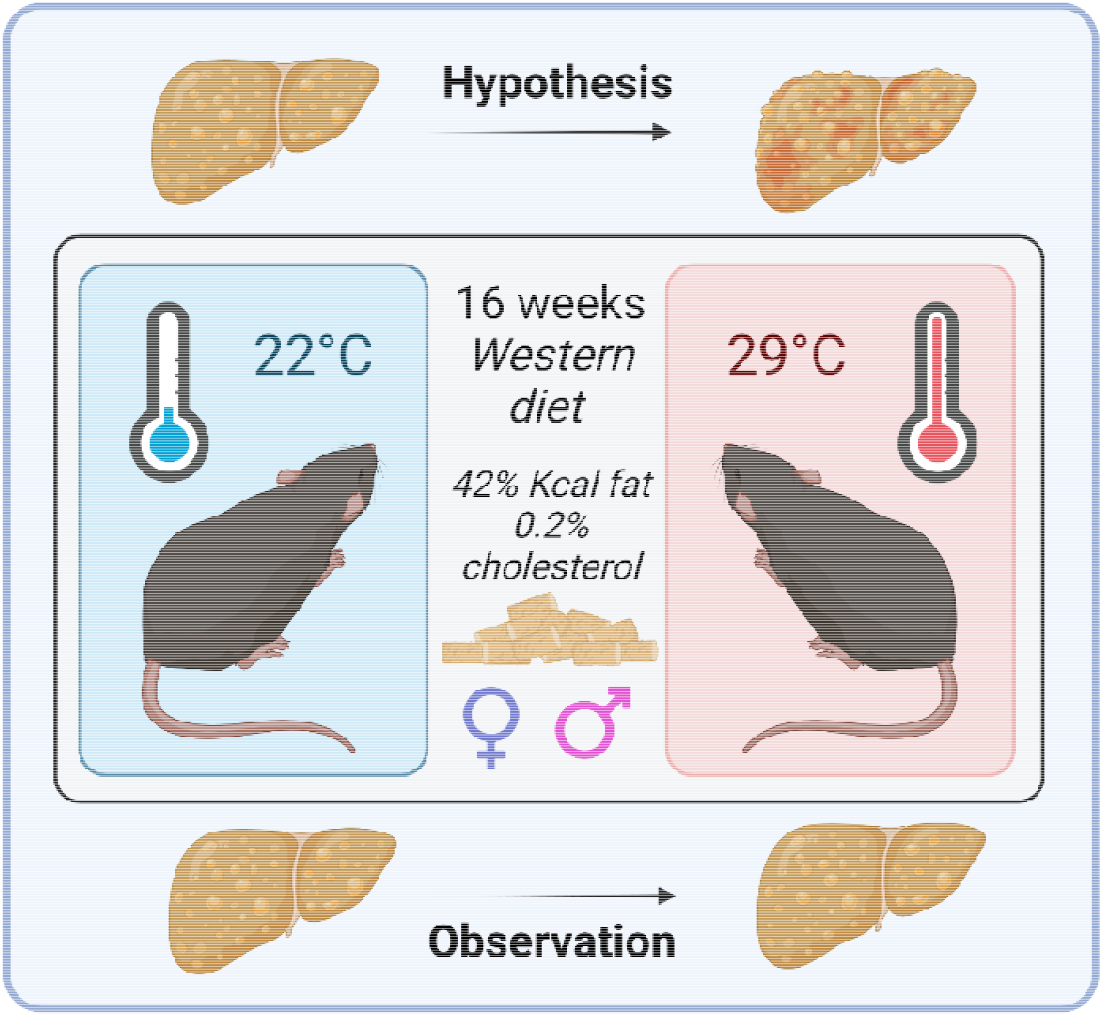

## Introduction

Metabolic dysfunction-associated fatty liver disease (MAFLD [1]) is the hepatic manifestation of obesity and metabolic syndrome. MAFLD is one of the largest global causes of chronic liver disease (with an unsettlingly high prevalence in children [2-4]) and is expected to affect over a third of adults by 2030 [5]. MAFLD encompasses a continuum of liver pathologies that begins with the benign steatosis in hepatocytes. In certain at-risk individuals, lipotoxic species can accumulate, leading to the onset of inflammation (non-alcoholic steatohepatitis; NASH) and fibrosis. Severe cases of NASH lead to hepatic cirrhosis or hepatocellular carcinoma [6-8]. Despite the identification of metabolic, fibrotic and inflammatory pathways for therapeutic targeting, there are currently no approved drugs to treat or halt progression of NASH, although clinical trials are ongoing [9]. Currently, only the early stages of benign steatosis are considered reversible with lifestyle changes and weight loss.

Critical to the effort to combat this chronic liver condition are the preclinical models used to interrogate potential targets. Rodent models of MAFLD are achieved via dietary intervention and/or genetic manipulation [10-12]. Dietary models include deficiency of choline and methionine first introduced in the 1930s and excess dietary fat being one of the most consistently altered components [13]. Genetic alterations consist of various naturally occurring human disease variants or mutations that alter metabolism such as leptin receptor mutations in obesogenic *ob/ob* mice. While these models achieve the phenotype of various MAFLD stages, the mechanisms by which the NASH-induced fibrosis are achieved are not always translationally relevant [14].

While diet and genetics have been at the forefront of controllable variables in mouse models, housing temperature has presented a new consideration. Conventional housing at 21-22°C induces cold stress in mice unlike the human population who, when clothed, are comfortable within this range [15]. This results in immunosuppression and brown adipose tissue-mediated thermogenesis in mice, two mechanisms that highly blunt NASH progression [16]. In recent years, numerous groups have highlighted the impact of housing temperature on different disease contexts. Giles *et al*. observed exacerbated fatty liver progression in high fat diet (HFD)- fed male and female mice housed at 30°C [17]. Mice housed at 30°C or thermoneutrality (T_N_) accumulated more fat mass due to lower BAT-mediated energy expenditure. Further, lower levels of corticosterone, an immunosuppressive stress hormone, exacerbated hepatic inflammation. Lastly, since female mice are more resistant to NASH development, the exacerbated phenotype observed at thermoneutrality highlights the importance of housing temperature [17]. This potential paradigm shift presents an opportunity to investigate key drivers of fatty liver progression that could better mimic human pathophysiology, but also calls into question whether past mechanistic insights hold true.

Mice fed a Western diet (WD) high in fat, sucrose and supplemented with a high dose of cholesterol, progress toward NASH after long periods (greater than 24 weeks) [18-20]. Accelerated timelines usually require supraphysiological concentrations of cholesterol (>0.2%) and/or genetic manipulations to achieve hepatic fibrosis [14; 21; 22]. Here we sought to address whether feeding a WD, high in fat, sucrose and with 0.2% cholesterol in conjunction with T_N_ housing could accelerate the progression toward NASH-induced fibrosis in C57Bl/6J male and female mice. While both sexes gained significantly more weight when fed a WD, only male mice housed under T_N_ conditions gained more weight compared to those at ambient room temperature (T_S_). There were select differences in the levels of circulating lipid and markers of hepatic injury and we observed an increase in the expression certain macrophage transcripts in both sexes. However, no differences in hepatic steatosis or pathological scoring were seen between WD-fed T_N_ and T_S_ mice. Finally, sex-specific and sex-independent effects were observed on markers of glucose homeostasis and insulin tolerance.

## Materials and Methods

### Mice

Wild-type mice were obtained from Jackson Laboratory and bred at the University of Ottawa, while housed at 22 °C. Male and female C57Bl/6J mice (12-15 weeks old) were acclimated for 2 weeks at either 22°C or 29°C in static cages. Mice were co-housed with littermates in groups of 2-5 mice. Once acclimated, mice were fed a 0.2% cholesterol high fat (41% Kcal) Western diet (Research diets – D12079B) or a macronutrient-matched low fat (10% Kcal) control diet (Research diets – D14020502) for 16 weeks. Mice were fasted for 4 hours prior to the tissue harvest. All animal protocols were approved by the uOttawa Animal Care Committee (BMIe3744).

### Non-terminal parameters

During week 15 of dietary intervention, mice underwent the following noninvasive procedures. Body composition was determined by ECHO MRI prior to the intraperitoneal glucose tolerance test (GTT). For the GTT, 4 h-fasted mice were injected with D-glucose (1.5 mg/kg *i*.*p*., made up in saline). For the insulin tolerance tests (ITT), 4 h-fasted mice were injected with insulin (0.8 U/kg *i*.*p*., made up in saline). Glucometer measurements for both tests were taken at baseline, 20-, 40-, 60-, 90- and 120-minutes post-injection. The above-mentioned tests were performed at 22°C to avoid acute effects of temperature on glucose clearance [23]. GTT and ITT were performed two days apart. For determination of intestinal permeability, mice were fasted for 6 hours followed by oral gavage of 150 µL of 80 mg/mL FITC-dextran (Sigma, 4 kDa). Blood was collected via tail vein at baseline and 1-hour post-gavage. Serum was collected via centrifugation and fluorescence measured by spectrophotometry (excitation 485 nM, emission 530 nM). Baseline serum readings were subtracted from 1-hour serum readings of each biological replicate. All samples were carefully protected from light.

### Serum and liver biochemical analysis

Terminal blood was collected via cardiac puncture of the left ventricle and placed on ice. Once clotted, serum was collected by centrifugation. Circulating insulin was determined by ELISA (Mouse Wide range insulin Elisa kit – Toronto BioScience Inc). Serum ALT, AST, ALP, TG, cholesterol, LDL and HDL were measured using the Beckman Coulter AU480 clinical chemistry analyzer in combination with appropriate reagents, calibrators and quality control materials. These readouts were performed by the Pathology core at Toronto Centre for Phenogenomics. Liver was snap frozen in liquid nitrogen and chipped on dry ice for Bligh and Dyer lipid extraction [24] and BCA protein quantification. Total liver cholesterol and triglycerides were determined with colorimetric assay kits (L-Type Triglyceride M – Fujifilm Wako, Infinity Cholesterol Liquid Stable Reagent – Thermo Fisher) and normalized to total protein content.

### Transcript expression

Liver was snap frozen in liquid nitrogen and chipped on dry ice. Frozen livers were homogenized in TriPure Isolation Reagent. Once equalized, RNA samples underwent reverse transcription (All-in-one RT Master Mix – ABM). qPCR was performed with BrightGreen Express reagent (ABM) and run on the CFX384 Real-Time PCR system for 40 cycles. Relative transcript expression was calculated using the delta-delta Ct method where genes of interest were normalized against the average Ct values of both *Actb* and *Tbp* [25], and shown relative to liver samples from male mice housed at T_S_ and fed a WD.

**Table 1.**
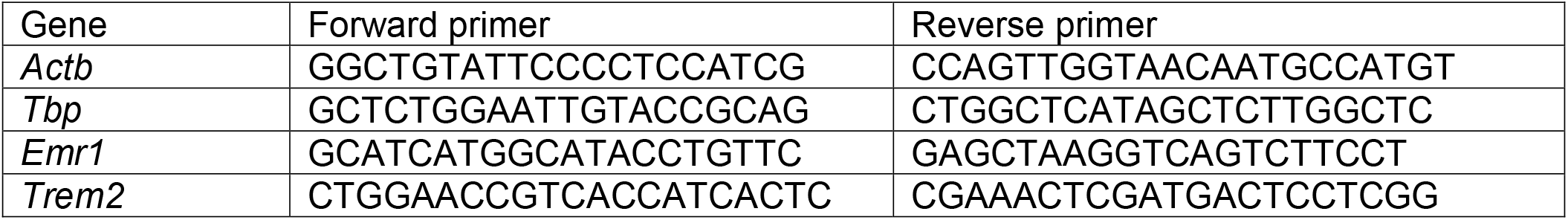
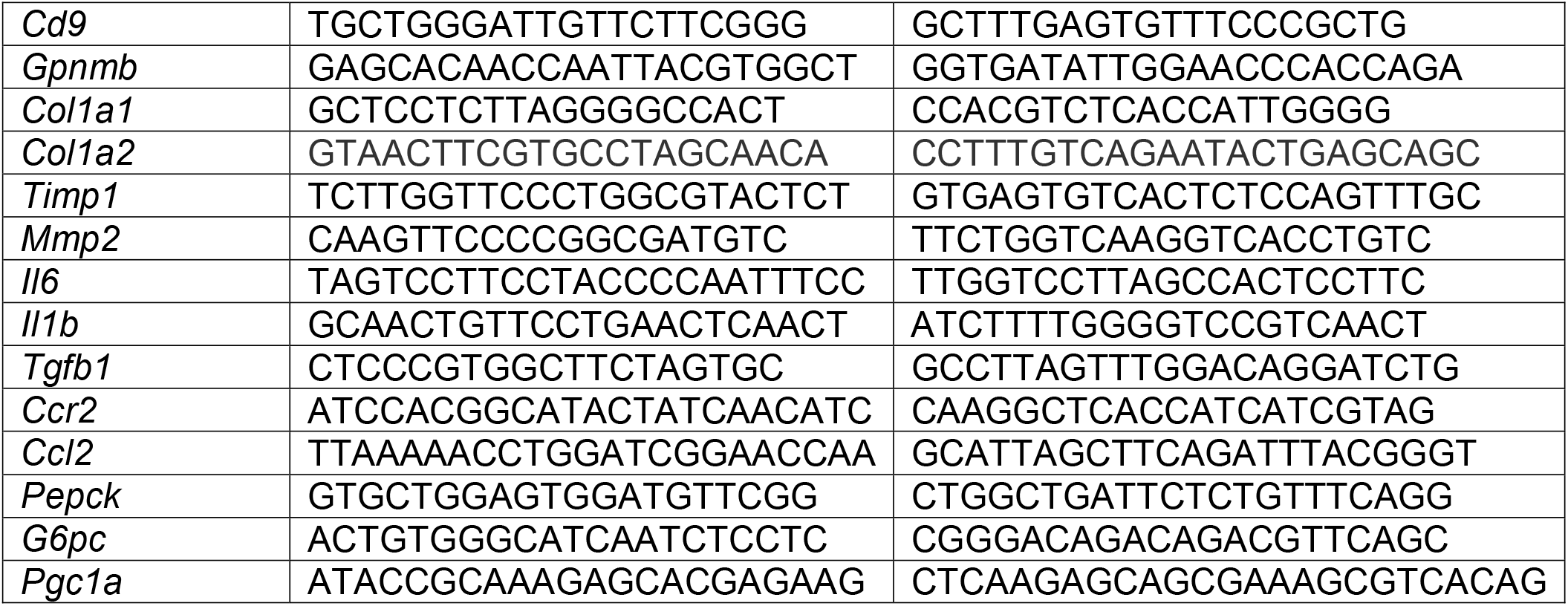
Primer sequences

### Histology

Following cardiac perfusion, liver tissue was fixed in 30 mL of 10% neutral buffered formalin. After 24 hours fixation, samples were placed in 70% ethanol and sent for processing at the University of Ottawa Louis Pelletier Histology Core Facility. Samples were paraffin-embedded, microtome sectioned and stained with H&E and Masson’s Trichrome. Slides were scanned at 20X magnification on the Axio Scan Z1 Slide Scanner. NAFLD score was qualitatively and blindly determined from H&E-stained sections by a third-party pathologist.

### Statistics

GraphPad Prism software was used for all statistical analyses. Two-way ANOVA was performed to compare between CD- and WD-fed mice housed under TN or TS conditions, where a Šídák multiple comparisons test was performed to determine significant differences between house temperature/within dietary treatment (figures 1, 2 and S1), between sexes/within housing temperatures (figure 3) and between housing temperature/within sex (figure 4). Comparisons between two groups (figure 1) were made using an unpaired, two-way Student’s t test. Data, where individual points are not displayed, represent mean ± SEM. A *p*-value of <0.05 was considered statistically significant.

**Figure 1.**
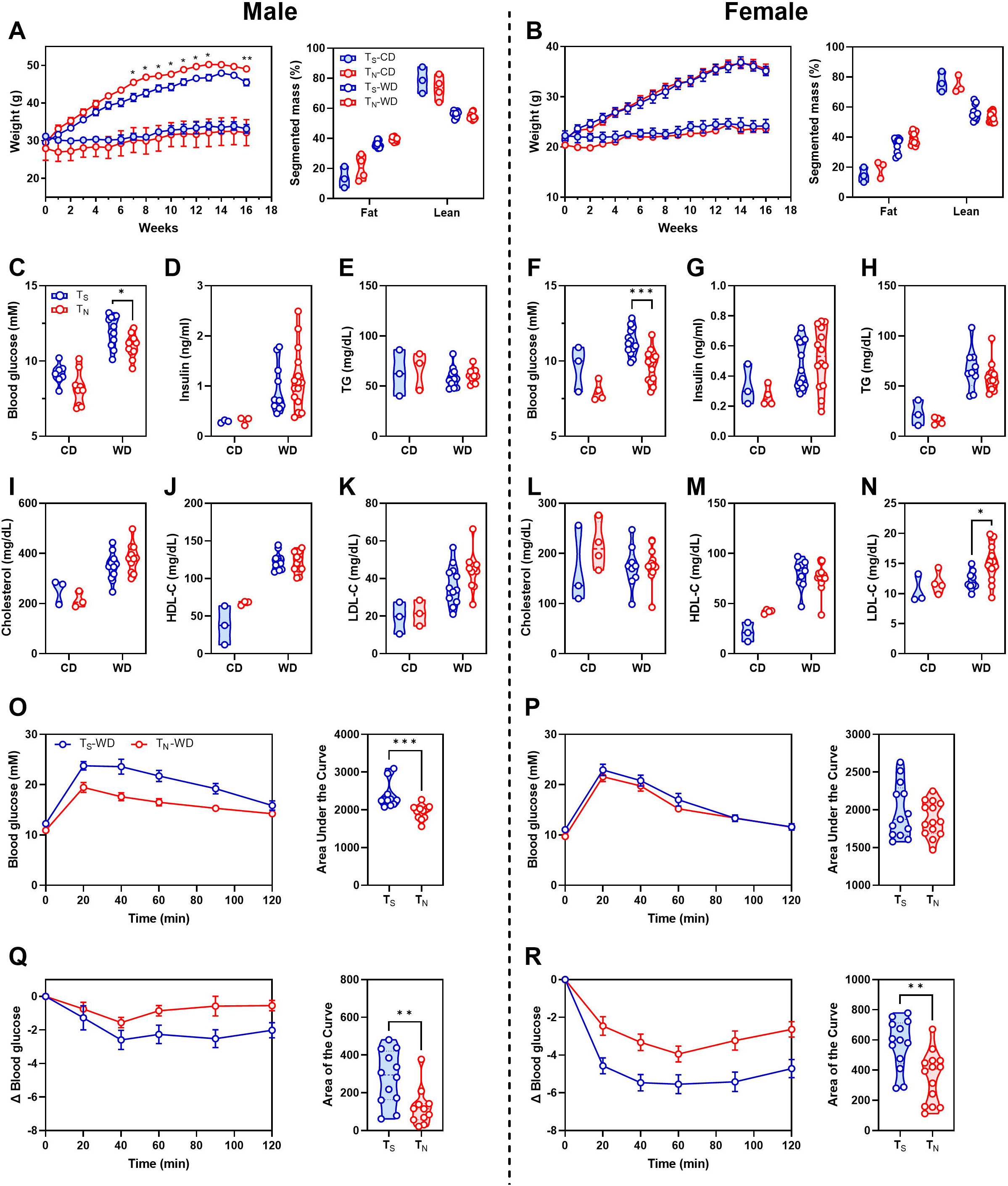
T_N_ housing and WD feeding has select effects in male and female mice. A/B) Male and female body weights, as well as the percentage of fat and lean mass for all groups, respectively. Mice were fasted for 4 h and blood was collected for serum determination of (for male and female mice, respectively) C/F) blood glucose, D/G) insulin, E/H) TG, I/L) total cholesterol, J/M) HDL-cholesterol, K/N) LDL-cholesterol. O/P) Male and female IPGTT of WD-fed mice in response to 1.5 g/kg glucose. Q/R) Male and female ITT of WD-fed mice in response to 0.8 U/kg insulin, where data is shown normalized to each individual animal’s baseline glucose level and the “area of the curve” is displayed. Data in A/B representing body weight are shown as mean ± SEM and were analyzed by a 2-way repeated measures ANOVA with Tukey test for multiple comparisons. Data shown in C-N were analyzed by 2-way ANOVA with a Šídák multiple comparisons test to determine significant differences within diet and between housing temperatures. Data in O-R are mean ± SEM where Student’s t test was used to determine differences between the two groups. Male and female CD groups were comprised of 3-4 mice, whereas male and female WD groups were comprised of 11-14 mice.

**Figure 2.**
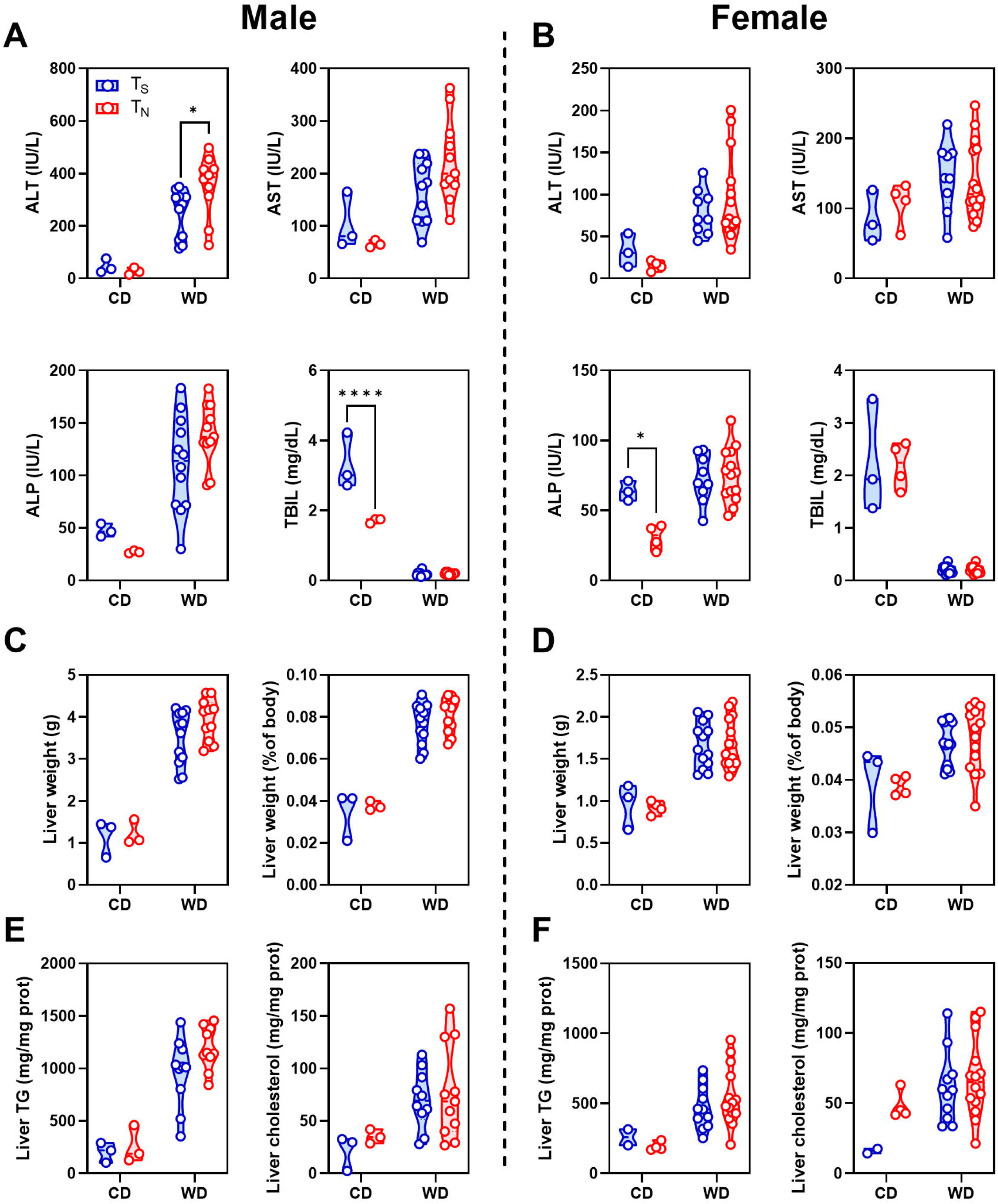
T_N_ housing and WD feeding does not alter liver enzyme or lipid levels. Serum determination of A) male and B) female alanine aminotransferase (ALT), aspartate aminotransferase (AST), alkaline phosphatase (ALP) and total bilirubin (TBIL) activity. C/D) Male/female liver weight (g) and as a percentage of the final body weight. E/F) Male/female liver TG and cholesterol levels. Data were analyzed by 2-way ANOVA with a Šídák multiple comparisons test to determine significant differences within diet and between housing temperatures. Male and female CD groups were comprised of 3-4 mice (although one liver from the female T_S_-CD group was lost in the tissue harvest, leaving n=2 for that group), whereas male and female WD groups were comprised of 11-14 mice.

**Figure 3.**
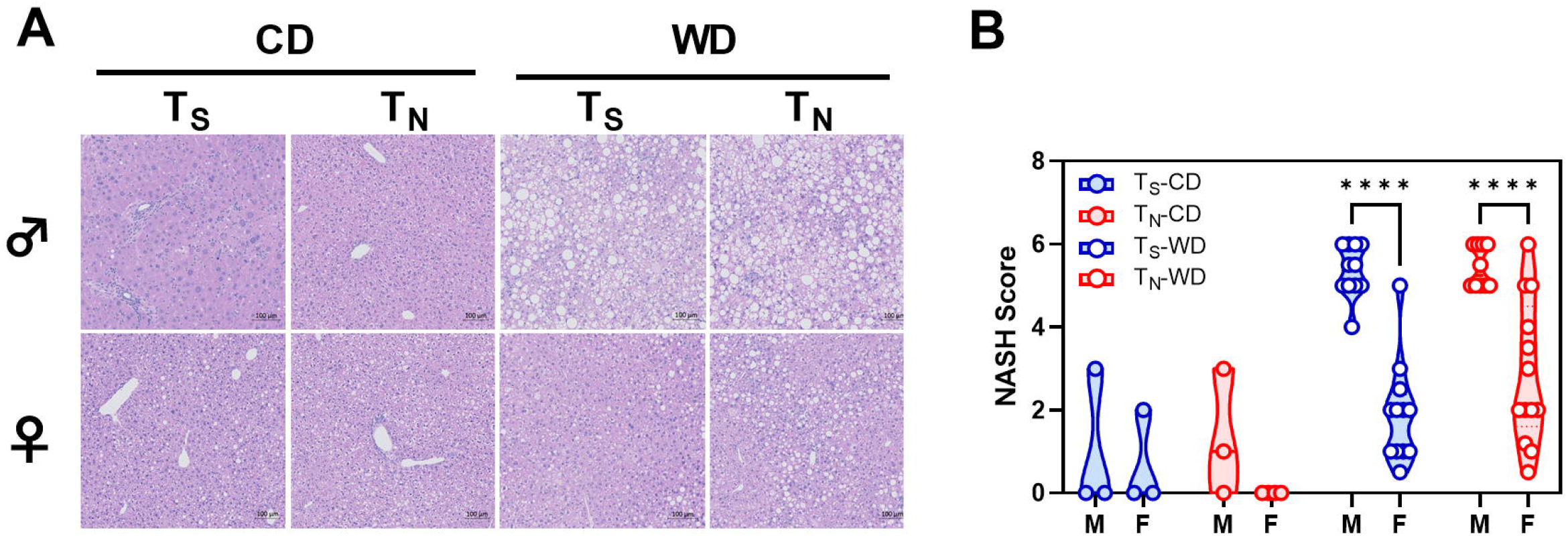
T_N_ housing does not exacerbate progression toward NASH beyond that observed with WD feeding. A) Representative H&E-stained images of liver taken from male and female mice (scale bar is 100 µm). B) NASH score representing histological scoring of steatosis, ballooning, inflammation and fibrosis. Data were analyzed by 2-way ANOVA with a Šídák multiple comparisons test to determine significant differences within diet and housing temperatures and between sexes. Male and female CD groups were comprised of 3-4 mice, whereas male and female WD groups were comprised of 11-14 mice.

**Figure 4.**
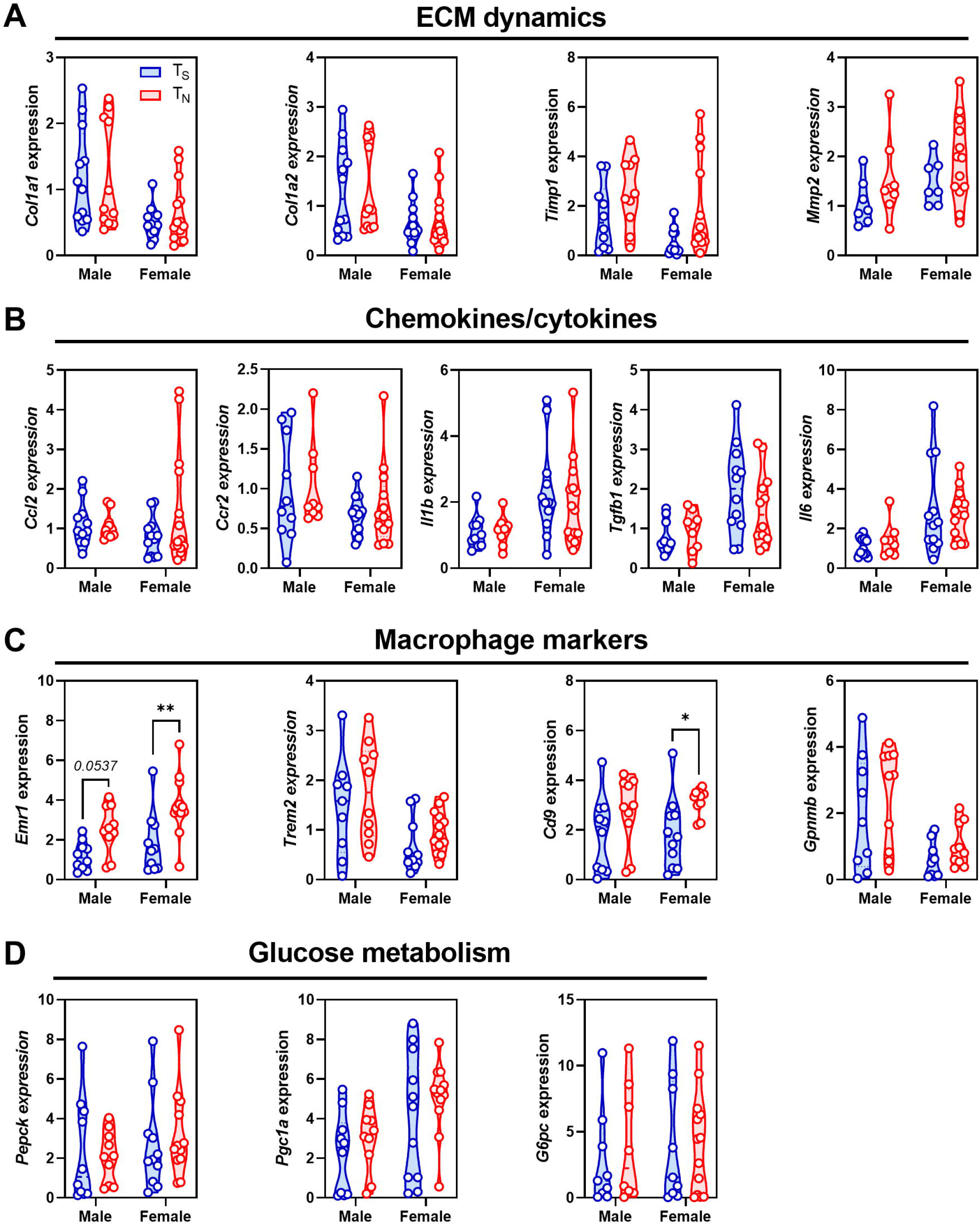
WD feeding has little effect on transcript markers of hepatic fibrosis, inflammation or gluconeogenesis in male and female mice housed at T_N_ or T_s_. Hepatic mRNA transcript levels from WD-fed male and female mice. A) markers of extracellular matrix (ECM)-regulating transcripts (*Col1a1, Col1a2, Timp1, Mmp2*), B) chemokine transcripts (*Ccl2, Ccr2*), C) cytokine transcripts (*Tgfb1, Il6, Il1b*), D) macrophage transcript markers (*Erm1, Cd9, Trem2, Gpnmb*), E) gluconeogenic transcripts (*Pepck, G6pc, Pgc1a*). Transcript expression was normalized to the average expression of *ActB* and *Tbp* and shown relative to male T_S_-WD-fed mice. Data were analyzed by 2-way ANOVA with a Šídák multiple comparisons test to determine significant differences within diet and between housing temperatures. Male and female WD groups were comprised of 6-14 mice.

## Results

### T_N_ housing and WD feeding affects body weight, glucose tolerance and insulin sensitivity differently in male and female mice

To evaluate the effectiveness of T_N_ housing on the progression of MAFLD, we used male and female C57Bl/6J mice and began WD or control diet (CD) feeding at ∼12 weeks of age. We acclimated T_N_-housed mice for 2 weeks prior to initiating the diet. Over the course of the 16-week intervention, males fed a WD and housed under T_N_ conditions gained significantly more weight compared to their T_S_ counterparts (**Figure 1A**). No differences were observed in males fed a CD at either housing temperature. Moreover, there were no differences in the percentage of lean or fat mass between WD-fed groups. No changes were observed in females with respect to weight gain or percentage lean and fat mass (**Figure 1B**). Following 16 weeks of dietary intervention, 5h-fasted blood glucose levels were consistently higher in WD-fed male and female mice compared to CD (**Table S1**). Interestingly, when observing the effects of housing temperature in WD-fed mice, blood glucose levels were lower in male and female mice kept at T_N_ (**Figure 1C and F**). While basal insulin levels were higher with WD feeding compared to CD (**Table S1**), there were no significant temperature-dependent differences between any of the groups in either sex (**Figure 1D and G**). While similar patterns were observed with circulating levels of triglyceride (TG), total cholesterol, HDL-cholesterol, LDL-cholesterol was significantly higher in WD-fed, T_N_-housed females (**Figure 1E and 1H-N**). Finally, due to the important role of the gut microbiome in MAFLD pathology, we assessed intestinal permeability to infer whether there could be increased hepatic exposure to pro-inflammatory intestinal microbial products. There were no differences between WD-fed mice housed at T_S_ or T_N_ conditions in circulating levels a fluorescently conjugated dextran substrate that was delivered via oral gavage (**Figure S1A**).

To assess the physiological response to glucose, we performed an intraperitoneal glucose and insulin tolerance test after 15 weeks of CD or WD feeding. No temperature-dependent differences were observed in CD-fed mice (**Figure S1B**). However, male but not female mice fed a WD and housed at T_N_ appeared significantly more glucose tolerant compared to their T_S_ counterparts (**Figure 1O and 1P**). To evaluate whether there were changes in sensitivity to insulin, mice were given an intraperitoneal dose of insulin based on total body weight. When blood glucose levels were plotted based on raw data, there were no significant differences between housing temperatures between either CD or WD-fed mice (**Figure S1B**). However, since there were significant differences in basal blood glucose levels, we analyzed the response to insulin within each individual animal by accounting for the initial glucose level of each mouse. Taking this into consideration, male and female WD-fed, T_N_-housed mice were less responsive to the glucose-lowering effect of insulin (**Figure 1Q and R**).

### Circulating liver enzymes and lipids are unchanged between T_N_ and T_S_ housing

The induction of hepatic steatosis and progression along the continuum to a NASH-like phenotype is typically associated with a decrease in liver function. As a proxy measure of liver injury, we determined the circulating levels of alanine transaminase (ALT), aspartate transaminase (AST), alkaline phosphatase (ALP) and total bilirubin (TBIL). As expected, WD feeding raised ALT, AST and ALP levels, and simultaneously reduced TBIL (**Table S1**). While ALT levels were further elevated in T_N_ males, there were no other differences in circulating markers when compared between temperature groups fed a WD (**Figure 2A-B**). The weight of the liver, both absolute and relative to body weight, was higher in WD-fed mice relative to CD (**Table S1**), independent of housing condition (**Figure 2C**). Similar results were observed in female mice, though to a lesser extent (**Figure 2D**). Total liver cholesterol and TG were not different between WD-fed T_S_ and T_N_ male or female mice, suggesting there were no changes in the degree of hepatic steatosis (**Figure 2E and F**).

### T_N_ housing and WD feeding does not render mice more susceptible to NASH progression, independent of sex

We next sought to qualitatively score the level of steatosis, hepatic ballooning, inflammation, and fibrosis as a pooled indicator of the progression toward NASH. Blinded assessment of H&E-stained histological sections showed that WD-fed male mice had significantly worsened progression toward NASH; however, contrary to our hypothesis, this was completely independent of whether the mice were housed at T_S_ or T_N_ (**Figure 3A, B**). We originally hypothesized that T_N_ housing would increase susceptibility to diet-induced hepatic dysfunction and cause female mice to progress toward a NASH-like phenotype more quickly; however, female mice remained significantly protected from NASH induction compared to male mice (**Figure 3A and B**). In addition to assessment of H&E staining, Masson’s trichrome displayed no collagen staining beyond the vascular regions in any of the WD-fed mice (**Figure S1C)**.

### T_N_ housing and WD feeding does not alter the expression profile of metabolic syndrome and NASH-related genes

To further validate and characterize the effect of WD feeding under T_s_ or T_N_ conditions, we determined the relative transcript levels of fibrotic (*Col1a1, Col1a2, Timp1, Mmp2*), macrophage (*Erm1, Trem2, Cd9, Gpnmb*), cytokine (*Il1b, Il6, Tgfb1*) and chemokine (*Ccl2, Ccr2*) markers in male and female mice. No significant differences in transcript expression related to extracellular matrix composition were observed between any of the groups (**Figure 4A**). Moreover, no differences were seen in cytokine or chemokine markers (**Figure 4B**). Specifically in female T_N_-WD, there was a significant increase in the transcript expression of *Emr1* and *Cd9*; however, there were no differences in the transcript expression of other lipid-associated macrophage (LAMs) subsets [26-28] (*Trem2, Gpnmb*) (**Figure 4C**). Finally, to follow up on the temperature-dependent differences in glucose metabolism, we assessed expression of genes involved in gluconeogenesis (*Pepck, G6pc, Pgc1a*) (**Figure 4D**) and observed no differences in expression.

## Discussion

Preclinical models have provided the foundation upon which much of our mechanistic understanding of fatty liver progression rests. Given the vast number of genetic and dietary/environmental instigators in mice, we sought to couple WD feeding with T_N_ housing conditions to potentially accelerate the onset and increase female susceptibility to diet-induced NASH. Contrary to our hypothesis, when C57Bl/6J mice were housed at T_N_ conditions and fed a WD for 16 weeks, there were no differences in markers of NASH-induced fibrosis progression or severity in either sex. Moreover, female mice remained partially protected against diet-induced hepatic dysfunction.

When exposed to a diet high in fat and housed at T_N_, mice of both sexes experienced a significant induction of immune-mediated hepatic dysfunction [17]. This high fat condition led to an increase in intestinal permeability and a microbe/TLR4/Th17-dependent response in male C57Bl/6 mice, independent of changes in body weight or adiposity. Male mice housed under T_N_ conditions and fed a WD were heavier than those housed at T_S_, which is entirely consistent with previous reports using a similar T_N_-WD protocol [29]. Despite this, there were no changes in intestinal permeability, markers of hepatic inflammation or fibrosis. In comparison to CD-fed male mice, WD-feeding augmented the transcript levels of markers of fibrosis, inflammation and glucose homeostasis; however, there were no changes due to housing temperature. Given the vast permutations of experimental conditions in the literature, it is extremely difficult to compare between studies. Many studies have used various dietary combinations containing high levels of fat and sugar (either as sucrose or fructose) and up to 2% cholesterol, for up to 1 year [22; 23; 30-34]. To our knowledge, the most similar design used a WD, which included 0.3% cholesterol as compared to 0.2% in our study, to demonstrate increased susceptibility of male C57Bl/6J mice to atherosclerosis following 24 wk of diet [29].Given that these mice are normally resistant to the initiation and progression of atherogenesis, these observations were attributed to increased inflammatory tone and elevated circulating cholesterol levels. No specific indicators of NASH progression were included.

There is a critical need to understand potential differences between how male and female mice progress toward fatty liver and NASH. While it is well known that female C57Bl/6 mice are more resistant to many of the effects of diet-induced obesity [35-37], we sought to directly compare the responses of male and female mice to T_N_ housing and WD feeding. Interestingly, T_N_ housing did not instigate further weight gain when paired with WD feeding in female mice, unlike their male counterparts and unlike high fat feeding [17]. However, there was a clear induction of weight gain in female mice fed a WD, independent of housing temperature and as reported previously [37]. Total and LDL-cholesterol rose significantly in WD-fed male, whereas in female levels were not different compared to CD, but significantly increased in response to T_N_ when fed a WD. Levels of HDL-cholesterol were similarly increased in both sexes with WD feeding. There were few unexpected differences in hepatic lipid levels and markers of liver health (**Figure 2**), with WD feeding augmenting levels in both sexes as previously reported [32], although ALT levels were higher in WD-fed males housed at T_N_ with a trend toward higher AST levels. Despite this, there were no significant changes in NASH scoring between housing temperatures. While female mice fed a WD and housed under T_S_ or T_N_ conditions were statistically indistinguishable, T_N_ females tended to have more severe scoring and had increased had higher expression of macrophage-specific markers (*Emr1* and *Cd9*), which was independent of differential effects on weight gain and adiposity. Recent studies have delineated multiple macrophage subsets in the hepatic MAFLD environment, including bone marrow-derived LAMs, defined in part by *Cd9* expression.

Prior studies using a range of high fat-containing diets have demonstrated that glucose tolerance tends to improve in male mice housed at T_N_ conditions [23; 30], although female data remains unavailable. Male mice fed a WD and housed at T_N_ were significantly more glucose tolerant compared to male mice housed at T_S_, while there were no differences between housing temperatures in females. Interestingly, while markers of liver lipid metabolism and inflammation were indistinguishable between T_S_ and T_N_-housed mice fed a WD, blood glucose levels after 5 h of fasting were significantly lower in both male and female mice (**Figure 1C, F**). Serum insulin levels after a comparable fast were also not different and there were no significant changes in the transcript expression of gluconeogenic genes in the liver, although these measures were increased by WD feeding, independent of temperature. When assessing insulin sensitivity, the impairment in insulin-associated drop in blood glucose was almost entirely mediated by the elevated baseline levels of glucose in T_S_ - compared to T_N_ -housed mice. This is best exemplified when the ITT values of each animal were expressed as a function of that same animal’s initial blood glucose and summarized as the area of the curve [38]. When C57Bl/6J mice were fed a diet high in fat (60% kcal from fat) and house at T_N_ conditions for 12 wk, there were no differences in glycemia or markers of glucose and insulin tolerance [23]. However, whether WD feeding, and T_N_ housing indeed impairs insulin action in peripheral or hepatic sites requires further study.

There remain important considerations when interpreting our results. Firstly, we defined T_N_ to be approximately 29°C (28.5-29.5°C) as opposed to 30°C. Currently there are discussions about the exact temperature of thermoneutrality, including whether this is a range or point [39]. Moreover, the strain of mouse must be considered [15; 40]. Given the diversity of approaches, comparisons between thermoneutral studies becomes challenging. Additionally, while cages were placed at either T_S_ or T_N_ conditions, mice were not singly housed. Despite the effect of huddling on elevating body temperature, mice are still considered cold stressed when co-housed at 22°C. Moreover, as most fatty liver/NASH studies co-housed littermate, we used this experimental paradigm to test our hypothesis. Co-housing was also meant to minimize the social anxiety that affects singly housed mice and to enable reciprocal fecal microbiota transfer between mice (coprophagy) [41; 42]. Secondly, it would have been informative to track food intake and markers of energy expenditure, as has been done previously [22; 43]. In male mice, T_N_ housing and WD feeding increased body weight, which may have stemmed from altered energy expenditure. However, this difference was inconsequential in the progression of hepatic steatosis and markers of NASH. Moreover, in female mice, where there was a tendency to show higher severity on the NASH score, this occurred without changes in weight. Finally, we identified that aspects of glucose homeostasis that were affected by WD feeding and T_N_ housing. However, though 16 wk was insufficient for T_N_ housing to exacerbate WD-induced progression toward NASH, we cannot rule out the potential for altered hepatic and/or circulating immune populations as well as transcriptional responses that may differentially prime the hepatic metabolic and immune landscape for progression toward NASH [33; 44].

T_N_ housing has the potential to validate and clarify molecular and metabolic mechanisms that underpin normal and pathophysiology. Our study revealed that WD feeding coupled with T_N_ housing does not exacerbate the progression of hepatic fibrosis and NASH-like pathologies, nor do female mice become significantly more susceptible. Future studies that integrate a longer treatment time, the potential inclusion of dietary cholesterol (which is known to proportionally instigate hepatic fibrosis and stellate cell activation in mice), as well as fructose supplementation (to exacerbate lipogenesis) in conjunction with thermoneutral housing will be necessary to provide a clearer translational model for interrogating mechanisms and testing potential therapies.

## Acknowledgments

The authors would like to thank the University of Ottawa Louise Pelletier Histology Core Facility for the processing, staining and scanning of the liver tissues, as well as Dr. Manijeh Daneshmand for pathological scoring of liver sections.

## Contributions

Author Contributions: J.R.C.N., T.K.T.S., P.G., C.O., N.A.T., H.D., C.P. and M.K.K. performed the experiments and testing. E.E.M. and M.D.F. provided funding. J.R.C.N. and M.D.F. wrote the manuscript. All authors edited the manuscript and provided comments. M.D.F. is the guarantor of this work and, as such, had full access to all the data in the study and takes responsibility for the integrity of the data and the accuracy of the analysis.

