## Supplemental material for "Thermoneutral housing does not accelerate metabolic dysfunction-associated fatty liver disease in male or female mice fed a Western diet"

### Figure S1

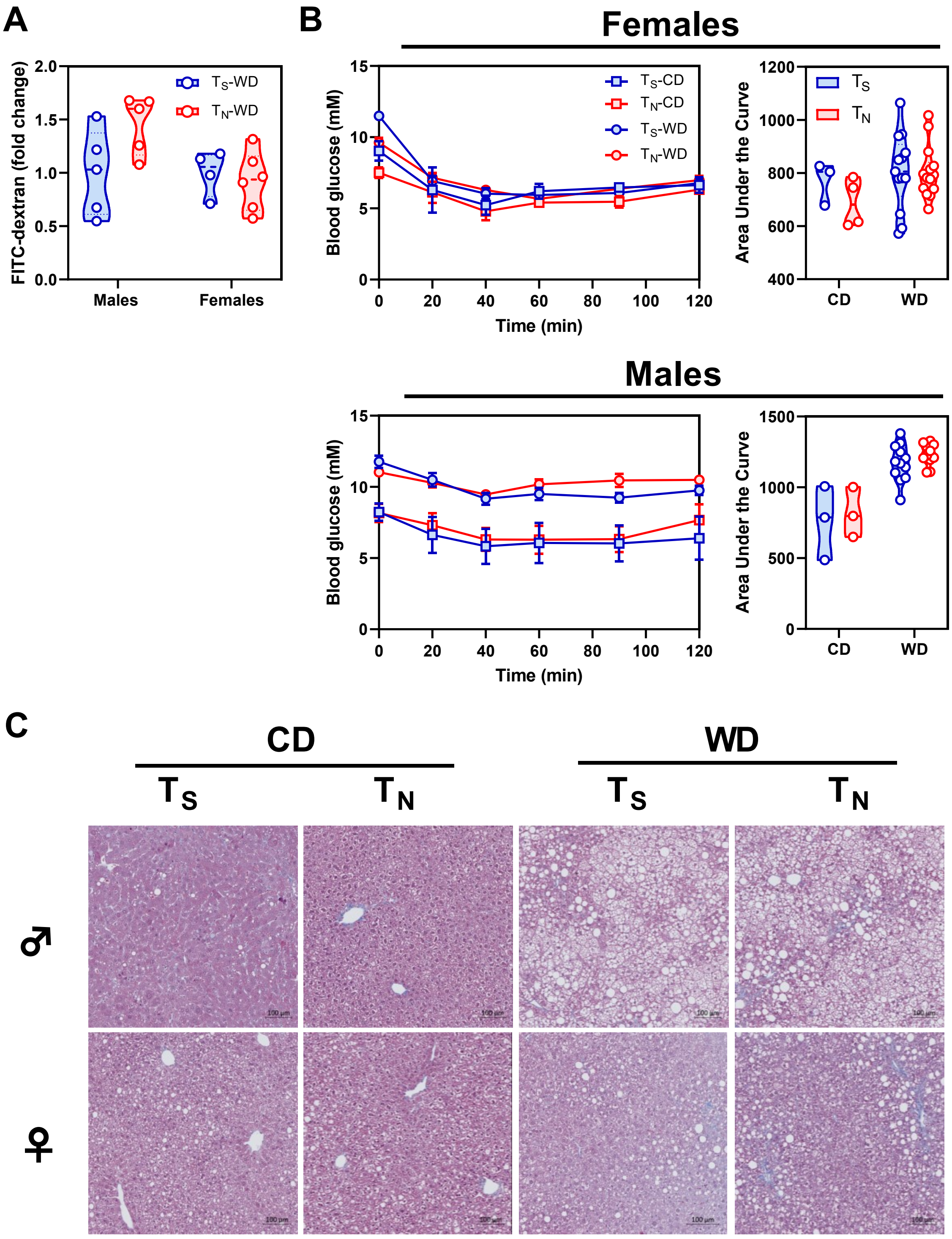
A
B
Females
Males
C
FITC-dextran (fold change)
Males
Females
TS-WD
TN-WD
Blood glucose (mM)
Time (min)
TS-CD
TN-CD
Area Under the Curve
TS
TN
CD
WD
♂
♀

**Supplemental Figure 1. $T_N$ housing and WD feeding effects on gut permeability, hepatic fibrosis and insulin sensitivity.** A) Male and female intestinal permeability assessment of WD-fed mice fluorescence of FITC-dextran in serum. B) Female and male ITT curves and area under the curve, shown as mM values. C) Representative Masson's Trichrome-stained images of liver of male and female mice (scale bar is 100 μm). Data in B representing ITT curves are shown as mean ± SEM. Data were analyzed by 2-way ANOVA with a Šídák multiple comparisons test to determine significant differences within diet and between housing temperatures. Data in A, male and female WD groups were comprised of 4-6 mice. Data in B, male and female CD groups were comprised of 3-4 mice, whereas male and female WD groups were comprised of 11-14 mice.

**Table S1. Statistical values of 2-way ANOVA multiple comparisons tests**

| | | **Male** | | | **Female** | | |
|---|---|---|---|---|---|---|---|
| **Measurement** | **Figure** | **Interaction** | **Diet** | **Temp** | **Interaction** | **Diet** | **Temp** |
| 5 h fasted Serum Glucose (mM) | 1C and 1F | 0.9611 | ***<0.0001*** | 0.0024 | 0.9805 | ***0.0005*** | ***0.0007*** |
| Random fed Serum Insulin (ng/ml) | 1D and 1G | 0.6863 | ***0.0044*** | 0.6427 | 0.4669 | ***0.0091*** | 0.874 |
| Serum TG (mg/dL) | 1E and 1H | 0.4623 | ***<0.0001*** | 0.2394 | 0.9259 | ***<0.0001*** | 0.2431 |
| Total Serum Cholesterol (mg/dL) | 1I and 1L | 0.156 | ***<0.0001*** | 0.9145 | 0.2383 | 0.3792 | 0.1735 |
| Serum HDL-C (mg/dL) | 1J and 1M | 0.0183 | ***<0.0001*** | ***0.0352*** | 0.0564 | ***<0.0001*** | 0.1148 |
| Serum LDL-C (mg/dL) | 1K and 1N | 0.5572 | ***0.0002*** | 0.278 | 0.4127 | ***0.0265*** | 0.0593 |
| Serum ALT (IU/L) | 2A and 2B | 0.1706 | ***<0.0001*** | 0.3304 | 0.3492 | ***0.0013*** | 0.9653 |
| Serum AST (IU/L) | 2A and 2B | 0.1092 | ***0.0013*** | 0.7053 | 0.5597 | ***0.0387*** | 0.715 |
| ALP (IU/L) | 2A and 2B | 0.1461 | ***<0.0001*** | 0.8109 | ***0.0222*** | ***0.0013*** | 0.0509 |
| TBIL (mg/dL) | 2A and 2B | ***<0.0001*** | ***<0.0001*** | ***<0.0001*** | 0.818 | ***<0.0001*** | 0.818 |
| Liver weight (g) | 2C and 2D | 0.4491 | ***<0.0001*** | 0.3255 | 0.7742 | ***<0.0001*** | 0.9387 |
| Liver weight (% of body weight) | 2C and 2D | 0.9069 | ***<0.0001*** | 0.3916 | 0.82 | ***0.0009*** | 0.9564 |
| Liver TG (mg/mg protein) | 2E and 2F | 0.4623 | ***<0.0001*** | 0.2394 | 0.3971 | ***0.002*** | 0.9495 |
| Liver cholesterol (mg/mg protein) | 2E and 2F | 0.8224 | ***0.0092*** | 0.5448 | 0.2837 | ***0.0106*** | 0.1104 |
